# Conjecture: Some auto-immune diseases are caused by the tissue, not a problem with the immune system

**DOI:** 10.1101/032300

**Authors:** Harlan Robins

## Abstract

Applying highly sensitive modern immune profiling techniques to the blood or inflamed tissue of auto-immune patients with one of the common T-cell mediated diseases fails to detect large clonal expansions or other signs of a dysregulated immune system. Additionally, these new methods have shown that self-reactive T-cells are found in the periphery (and are not completely negatively selected in the thymus as previously believed). Combining these new data with well-established studies in auto-immunity leads to the conjecture that certain auto-immune diseases are likely to be caused by defects in tissue, not by dysregulation of the adaptive immune system. In particular, one hypothesis is that specific tissues up-regulate HLA class 2 genes, presenting epitopes that are bound by CD4^+^ helper T cells, which facilitates an immune response against the tissue specific cells.

---

The ability to profile the adaptive immune system at high throughput is leading to an unprecedented understanding of immune health. This knowledge is challenging our interpretation of the classical T-cell mediated autoimmune diseases. Now that we are confident in our ability to detect perturbations and deficiencies in the adaptive immune system, we can no longer attribute our failure to detect such alterations in studies of autoimmune disease to sensitivity limitations. We have deeply sequenced the T-cell receptors (TCRs) from clones in the blood in several autoimmune disorders, and we have sequenced TCRs from CSF in multiple sclerosis (MS) patients, colons of Crohn’s patients, pancreases of Type 1 diabetes (T1D) patients, and synovial fluid in Rheumatoid Arthritis (RA) patients. In none of these cases did we observe excessive clonal expansions in the cellular adaptive immune system. The changes we do observe appear no more drastic than a typical immune response to a pathogen, indicating that, with only a few exceptions (e.g. ALPS), the underlying cause of typical autoimmune diseases is likely not a defect in excessive clonal T-cell expansion or broad dysregulation of the cellular adaptive immune system.

In addition to this, the association between several autoimmune diseases and specific HLA class 2 alleles is well-established (*1*). For example, different HLA-DR-DQ alleles have been associated with extremely high hazard ratios for the onset of T1D, MS, and RA.

Combining these observations with new data coming from the lab of Mark Davis at Stanford suggests a different cause of autoimmune disease than what has been previously considered. In Mark Davis’ recent work (*2*), he showed that the principle of negative selection of T cells in the thymus is not as absolute as previously believed (*3*). Although thymic negative selection does play an important role in T-cell tolerance (*4, 5*), there is mounting evidence contradicting the dogma that all T-cells that bind to self-antigens are deleted. Although the mechanisms by which self-tolerance is maintained in the presence of autoreactive T-cells circulating in the periphery is not completely understood, it likely involves CD4^+^ T regulatory cells (*6*). Davis hypothesizes that these cells are not deleted because the whole complement of T-cells -including those that bind to self-antigens-need to be present in order to prevent ‘holes’ in the immune system that could lead to lowered ability to identify pathogens. Jenkins and colleagues have recently shown that indeed some TCRs that escape negative selection are cross-reactive to both self- and pathogen-derived antigens (*7*), demonstrating that autoreactive T-cells may play an important role in protecting the host from some pathogens.

Together, these data lead me to conjecture that the classic T-cell mediated autoimmune diseases (such as MS, T1D, Crohn’s disease, RA, etc.) are not always caused by an immunological defect. Instead, I propose that in some cases, they are caused by a defect in the relevant tissue, and that this defect potentially manifests as the aberrant expression of the normally silent HLA class 2 genes in non-Antigen Presenting cells (nonAPCs), and the ensuing surface-presentation of self epitopes. As a consequence of this, CD4^+^ helper T-cells would bind these aberrant nonAPC cells and become activated, thus eliciting the tissue-specific chronic inflammatory injuries that are the hallmark of autoimmunity.

Gene expression control is mediated in a tissue-specific manner through several epigenetic mechanisms, including promoter binding by transcription factors, DNA methylation, and histone deacetylation. The expression of HLA class 2 genes has been observed in nonAPC tissue cells that break tolerance(*8, 9*). In addition, there is evidence from Lindsey Criswell’s group and others that different class 2 alleles have varied methylation patterns in certain cell types, which could lead to differential expression.

The HLA-DR-DQ alleles associated with T1D, MS, and RA have been shown to have different epigenetic markers in some tissues, although most studies have been limited to T-cells. Additionally, these alleles could be de-repressed by a variety of additional mechanisms due to the differences in their nucleic acid primary sequences. Since gene expression patterns are cell-type specific, the modification of these patterns will also be tissue-specific. I hypothesize that these changes act to turn on the HLA-DR-DQ genes (and possibly other class 2 genes) in specific tissues affected in a particular autoimmune disease. For example, the allele HLA-DRB1*04:01 could cause the locus to become expressed in beta islet cells for T1D patients (*10*). In turn, these beta islet cells would present the HLA class 2/peptide complexes, initiating direct CD4^+^ T-cell cytotoxicity and/or CD4^+^ T helper responses. The CD8^+^ killer T-cells would then have the help they need to kill the specific target cells, which are presenting HLA class 2 epitopes (in the case of T1D, these would correspond to beta islet cells). In principle, any gene(s) from the tissue could be involved in the immune response. In practice, a particular set of genes is more likely to be involved. For example, gene X might code for a really strong class 2 epitope that is presented by HLA-DRB1*0401 proteins or some strong class 1 epitopes. Then, when the islet cells express gene X, the CD8^+^ T-cells will attack, with the corresponding CD4^+^ T help needed.

This hypothesis addresses a set of confounding issues in autoimmunity. The tissue specificity of each disease is an open question. Why does an allele in HLA-DRB1*0401 cause an autoimmune response in beta islet cells? This specificity would suggest a tissue-specific cause, not a systemic immune dysfunction. There are many lines of evidence showing that autoreactive T-cells target tissue-restricted antigens (such as insulin or MOG) to mediate disease. However, in some cases this tissue-specific autoimmune attack is mediated by T-cells specific for ubiquitously-expressed antigens. For example, HSP60, HSP70, GAD65, and ICA69 are all expressed in multiple tissues including pancreatic beta cells, and are all known targets of autoimmune T-cell attack in T1D(*11*). Why do T-cells specific for these antigens attack beta cells in the pancreas of T1 diabetics and not other tissues?

Unfortunately, this hypothesis suggests that present treatment modalities will have limited success because they focus on treating the immune response, and not the tissue, which is the proposed cause of the autoimmune disease. Thus, if this hypothesis (which requires testing) is correct, it predicts that treatments that have shown early success such as bone marrow transplants to treat MS are might be temporary for some patients. On the other hand, perhaps a new class of therapies that treat the affected tissue could be developed.

## Acknowledgements

HR thanks Gerald Nepom, Ruth Tanaguchi, Mark Klinger, Marissa Vignali, Ryan Emerson, and Anna Sherwood for providing expertise in autoimmune disease and improving this hypothesis.

## References

1. G. T. Nepom, H. Erlich, MHC class-II molecules and autoimmunity. Annu Rev Immunol 9, 493–525 (1991).

2. W. Yu et al., Clonal Deletion Prunes but Does Not Eliminate Self-Specific alphabeta CD8(+) T Lymphocytes. Immunity 42, 929–941 (2015).

3. Y. Xing, K. A. Hogquist, T-cell tolerance: central and peripheral. Cold Spring Harb Perspect Biol 4, (2012).

4. M. S. Anderson et al., Projection of an immunological self shadow within the thymus by the aire protein. Science 298, 1395–1401 (2002).

5. K. Nagamine et al., Positional cloning of the APECED gene. Nat Genet 17, 393–398 (1997).

6. E. M. Shevach, Regulatory T cells in autoimmmunity*. Annu Rev Immunol 18, 423–449 (2000).

7. R. W. Nelson et al., T cell receptor cross-reactivity between similar foreign and self peptides influences naive cell population size and autoimmunity. Immunity 42, 95–107 (2015).

8. G. F. Bottazzo et al., In situ characterization of autoimmune phenomena and expression of HLA molecules in the pancreas in diabetic insulitis. N Engl J Med 313, 353–360 (1985).

9. U. Walter et al., Pancreatic NOD beta cells express MHC class II protein and the frequency of I-A(g7) mRNA-expressing beta cells strongly increases during progression to autoimmune diabetes. Diabetologia 46,1106–1114 (2003).

10. J. A. Noble, H. A. Erlich, Genetics of type 1 diabetes. Cold Spring Harb Perspect Med 2, a007732 (2012).

11. B. O. Roep, M. Peakman, Antigen targets of type 1 diabetes autoimmunity. Cold Spring Harb Perspect Med 2, a007781 (2012).

